# Origin and Evolution of DNA methyltransferases (DNMT) along the tree of life: A multi-genome survey

**DOI:** 10.1101/2020.04.09.033167

**Authors:** Madhumita Bhattacharyya, Subhajyoti De, Saikat Chakrabarti

## Abstract

**Background:** Cytosine methylation is a common DNA modification found in most eukaryotic organisms including plants, animals, and fungi. (Cytosine-5)-DNA methyltransferases (C5-DNA MTases) belong to the DNMT family of enzymes that catalyze the transfer of a methyl group from S-adenosyl methionine (SAM) to cytosine residues of DNA. In mammals, four members of the DNMT family have been reported: DNMT1, DNMT3a, DNMT3b and DNMT3L, but only DNMT1, DNMT3a and DNMT3b possess methyltransferase activity. There have been many reports about the methylation landscape in different organisms yet there is no systematic report of how the enzyme DNA (C5) methyltransferases have evolved in different organisms.

**Result:** DNA methyltransferases are found to be present in all three domains of life. However, significant variability has been observed in length, copy number and sequence identity when compared across kingdoms. Sequence conservation is greatly increased in invertebrates and vertebrates compared to other groups. Similarly, sequence length has been found to be increased while domain lengths remain more or less conserved. Vertebrates are also found to be associated with more conserved DNMT domains. Finally, comparison between single nucleotide polymorphisms (SNPs) prevailing in human populations and evolutionary changes in DNMT vertebrate alignment revealed that most of the SNPs were conserved in vertebrates.

**Conclusion:** The sequences (including the catalytic domain and motifs) and structure of the DNMT enzymes have been evolved greatly from bacteria to vertebrates with a steady increase in complexity and specificity. This study provides a systematic report of the evolution of DNA methyltransferase enzyme across different lineages of tree of life.

## Background

The genomes of eukaryotes are marked with regionally restricted epigenetic information responsible for regulating local activity states. Most widely studied epigenetic modification in humans is cytosine methylation which occurs almost exclusively in the context of CpG dinucleotide. The CpG dinucleotides tend to cluster in regions called CpG islands (Bird A. et. al. 2002). About 60% of human gene promoters are associated with CpG islands and are usually unmethylated in normal cells while some of them (~6%) become methylated in a tissue-specific manner during early development or in differentiated tissues (Brown and Strathdee, 2002). In general, CpG-island methylation is associated with gene silencing, genomic imprinting, X chromosome inactivation in females, histone modification, chromatin remodeling etc. DNA methylation and DNA methylation–associated proteins not only participate in gene transcription regulation in *cis,* but also act in *trans,* being involved in nuclear organization and in the establishment of specific chromosomal territories. Hypermethylated CpGs are needed to protect chromosomal integrity, which is achieved by preventing reactivation of endoparasitic sequences that cause chromosomal instability, translocations and gene disruption (Okano M. et. al. 1999).

DNA methylations in mammalian systems are observed throughout the genome barring the CpG islands (CGIs) (Bird A. 2002, Bird A. P. 1986). In fungi that have genomic 5-methylcytosine (m5C), only repetitive DNA sequences are methylated (Selker E. U. et. al, 2003). The most frequent pattern observed in invertebrates is ‘mosaic methylation’, comprising domains of heavily methylated DNA interspersed with domains that are methylation free (Bird A. P. et. al. 1979, Tweedie S. et. al. 1997). The highest levels of DNA methylation among all eukaryotes have been observed in plants, with up to 50% of cytosine being methylated in some species (Montero L. M. et. al, 1992). In maize, for example, such high levels seem to be due to large numbers of transposons, the degenerate relics of which dominate inter-genic regions and are targeted for methylation (Palmer et. al. 2003, SanMiguel P. et. al. 1996). However, other plants, such as *Arabidopsis thaliana,* display a mosaic DNA methylation pattern that is reminiscent of invertebrate animals.

Although DNA methylation appears to be a widespread epigenetic regulatory mechanism, genomes are methylated in diverse ways in different organisms. In animals, DNA methylation occurs mostly symmetrically (both strands) at the cytosines of a CG dinucleotide. DNA methylation in plant genomes can occur symmetrically at cytosines in both CG and CHG (H = A, T, or C) contexts, and also asymmetrically in a CHH context, with the latter directed and maintained by small RNAs (Law and Jacobson, 2010). In the model plant *Arabidopsis thaliana,* levels of cytosine methylation at CG, CHG, and CHH nucleotides are about 24%, 6.7%, and 1.7%, respectively (Cokus S. J. et. al. 2008, Lister R. et. al. 2008). However, it is important to bear in mind that the global DNA methylation pattern seen in vertebrates is by no means ubiquitous among eukaryotes. Several well-studied model systems have no recognizable *DNMT* like genes and are devoid of DNA methylation (for example, the yeast *Saccharomyces cerevisiae,* fruit fly *Drosophilla melanogaster* and the nematode worm *Caenorhabditis elegans).*

Despite the different methylation sequence contexts, cytosine methylation is established and maintained by a family of conserved DNA methyltransferases (Goll and Bestor 2005, Chan S. W. et. al 2005, Cheng and Blumenthal 2008). Not surprisingly, the absence of DNA methylation in some eukaryotes such as yeast, fruit fly, and roundworm is associated with the evolutionary loss of DNA methyltransferase homologs (Goll and Bestor 2005). Different C5-cytosine methyltransferases have been characterized in prokaryotes and eukaryotes. All methyltransferases share a catalytic domain containing 10 conserved small motifs, suggesting a common origin (Posfai J. et al. 1989; Kumar S. et al. 1994). Based on sequence similarity, the eukaryotic methyltransferase have been grouped into different subfamilies at different times, based on different criteria. For example, methylases were classified based of the methylation residue (M4C, M6A, and M5C), methylation activity *(“de novo* methylation” or “maintenance methylation”), and methylation state of substrate DNA (unmethylated / hemimethylated).

M6A and M4C methyltransferases are responsible for methylation of adenine residue (at N6) and cytosine residue (at N4), respectively. Both of these enzymes are primarily found in prokaryotes where it functions as restriction endonuclease when DNA is unmethylated and functions as methyltransferase when DNA is hemimethylated. M5C methyltransferases are responsible for methylation in cytosine residue (at N5) position and are found to have a large C-terminal catalytic domain and smaller N-terminal recognition domain. Until recently only one DNA methyltransferase, *DNMT1,* had been cloned from human and mouse cells. Recently, another group of DNA methyltransferases, *DNMT3A* and *DNMT3B* was isolated by database search.

DNMT1 is the most abundant DNA methyltransferase in mammalian cells, and considered to be the key maintenance methyltransferase in mammals (Okano M. et. al. 1999). The carboxyterminal catalytic domain of DNMT1 is responsible for transfer of methyl groups from S-adenosyl methionine to cytosines in CpG dinucleotides. The longer N-terminal portion of DNMT1 contains regions responsible for targeting the replication foci. A cysteine-rich Zn-binding motif and a polybromo domain that resemble regions of proteins known to have roles in chromatin modification (Leonhardt H et. al. 1992, Chuang L. S. et. al. 1997) are utilized for discriminating between unmethylated and hemi-methylated DNA, DNMT3 is a family of DNA methyltransferases that could methylate hemimethylated and unmethylated CpG at the same rate. The architecture of DNMT3 enzymes is similar to that of DNMT1, with a regulatory region attached to the catalytic domain. There are three known members of the DNMT3 family: 3A, 3B, and 3L. DNMT3A and DNMT3B can mediate methylation-independent gene repression. DNMT3A can co-localize with heterochromatin protein (HP1) and methyl-CpG-binding protein (MeCBP). They can also interact with DNMT1, which might be a co-operative event during DNA methylation. DNMT3A prefers CpG methylation to CpA, CpT, and CpC methylation, though there appears to be some sequence preference of methylation for DNMT3A and DNMT3B. DNMT3L contains DNA methyltransferase motifs and is required for establishing maternal genomic imprints, despite being catalytically inactive. DNMT3L is expressed during gametogenesis when genomic imprinting takes place. The loss of DNMT3L leads to bi-allelic expression of genes normally not expressed by the maternal allele. DNMT3L interacts with DNMT3A and DNMT3B and colocalizes in the nucleus (Okano M. et. al. 1999).

All reported DNMTs have a well conserved C terminal catalytic domain containing an ordered set of sequence motifs which alternates with non-conserved regions. Depending on the criteria used to define the limits of the conserved blocks, up to ten motifs can be identified (Lauster R. et. al. 1989, Posfai J. et. al. 1989, Klimasauskas S. et. al. 1989). In the original analysis of 13 M5C-methyltransferases, five motifs were considered highly conserved (I, IV, VI, VIII, and X), and the remaining five moderately conserved. Analysis of 36 sequences resulted in the inclusion of a sixth motif, motif IX, within the highly conserved set (Cheng X. et. al. 1993).

Different methyltransferases were reported in different organism and few attempts were made to explore their origin and evolution. Based on these observations, Colot and Rossignol (1999) proposed that methylation has divergent functions in different organisms, consistent with the notion that it is a dynamically evolving mechanism that can be adapted to perform various functions. Another report was published in 2005 where Ponger and Li tried to analyze the variability of methylation systems, with a survey of methytransferases in complete or almost complete eukaryotic genomes, including several species of Protozoa. They also reconstructed a phylogeny of the putative enzymes identified to study the evolutionary history of this function and to classify eukaryotic methyltransferases (Ponger and Li, 2005). Functional and structural conservation of DNMTs in human, mouse and cattle were observed along with similar patterns of transcript abundance for all of the proteins at different stages of early embryo development. Greater degree of structural similarity between human and bovine was observed for all of the DNMT (DNMT1, DNMT3A, DNMT3B, and DNMT3L) than that between human and mouse (Rodriguez-Osorio N. et. al. 2010).

Using the information provided by National Center for Biotechnology Information (NCBI) taxonomy, a phylogenetic tree was reported in a science perspective where methylation extent and type of methyltransferases were put together. The report suggested that the last common ancestor of eukaryotes contained a functional DNA methylation system with secondary expansion and the loss of methylation in some lineages while primitive methylation likely occurred at low to intermediate levels and was targeted to gene bodies and transposable elements, leaving gene promoters unmethylated (Jeltsch A, 2010). Zemach et. al. reported a quantitative estimation of DNA methylation in 17 species and found out that gene body methylation is conserved on the contrary to selective transposon methylation (Zemach A. et. al 2010). Another publication from the same group reported transposable elements and sex to be the major forces driving the evolution of methylation (Zemach and Zilberman, 2010). Shotgun bisulfite sequencing (BS-seq) was used to compare DNA methylation in eight diverse plant and animals. Different patterns of methylation were detected in flowering plants and animals (Feng S. et. al 2010). In one of the largest phylogenetic analysis till date comprising of 2300 sequences the evolutionary relationship among different DNMT1 and DNMT3 was explored. This study proposed a consensus model of the phylogeny of DNA methyltransferases indicating that DNMT1 and DNMT3A/B enzymes have an independent origin in the prokaryotic DNA methyltransferase sequence space and all were derived from methyltransferases of restriction modification systems (Jurkowska and Jeltsch, 2011).

DNA methylation machinery was always a central question of interest and a good number of reports are available about identification and diversification of DNA methylation machinery in higher order organisms and/or having small number of representatives from different taxonomical groups. However, a comprehensive overview of modifications in sequence and structural level over time and taxonomical hierarchy is still lacking. This study attempts to make a detail investigation of evolution of DNA methyltransferase enzymes along all branches of tree of life (TOL) including as many possible organisms from lower (bacteria) to higher (human) order of taxonomy. The objectives of this work are to search and identification of the homologues for DNA methyltransferase in all available genomes followed by a thorough and systematic comparison of the sequences in order to elucidate the evolution of DNMTs along the lineages of tree of life. Finally, naturally occurring variations were compared with SNPs observed in human populations to investigate if the evolutionary selection pressure on structural and functional motifs of DNMT was similar in genetic variations observed in human populations.

To reconstruct the tree of life based on DNA methyltransferase using phylogenetic methods, sequences were classified systematically from seven kingdoms. Parameters like gene copy number, sequence length and identity were compared across the kingdoms. Functional motifs and domains were identified and compared across different lineages of the tree of life. Conservation of each site/residue was measured along each kingdom group and finally compared with the available SNP data. All three groups of sequences were found to have a clear evolutionary pattern across kingdoms. Kingdom specific conserved functional motifs and domains were also observed. Moreover, to the best of our knowledge, this is the first large scale systematic report of evolution of DNMT enzymes across different lineages of tree of life including all possible organisms. A clear pattern of evolution was observed where the sequences of the enzyme achieved more complexity and specificity in higher order organisms. Comparison with single nucleotide polymorphism data in human shows that none of the SNPs were overlapping with functional motifs though more than 60% are conserved residues, while more than 80% SNPs occurring in non-conserved residues are tolerated.

## Results

### Collection and Classification of DNMT sequences

Cytosine (5) methyltransferase domain (Pfam ID: PF00145) was identified from Pfam database (Punta M. et. al. 2010) and was scanned to search similar sequences against different sequence databases like NCBI non redundant database (Coordinator N R 2018) and Uniprot (Leinonen R. et. al. 2004). Total 4845 unique sequences were identified in 710 organisms from all three databases.

For each type of DNA methyltransferases, (namely DNMT1, DNMT3A, DNMT3B and DNMTL) full length, annotated and experimentally reviewed sequences were collected from Uniprot (Leinonen R. et. al. 2004). 55 DNMT1 unique sequences were identified whereas 17, 26 and 5 unique sequences were extracted for DNMT3A, DNMT3B and DNMTL, respectively. For each type of DNMT enzyme a hidden markov model (HMM) profile was created **(See Method)** and each of the collected 4845 sequences was scanned against these signature DNMT profiles. Finally each sequence was classified to a particular DNMT group based on its alignment matching with the respective DNMT HMM profile. From the initial 4845 sequences, 1783 sequence were found to be DNMT1, whereas only 67 and 172 sequences were identified to be DNMT3A and DNMT3B, respectively. No DNMTL sequences were identified. The protocol for sequence identification is described as a flow chart in **Figure S1**.

In this study 16S rRNA based tree of life (Yarza P. et. al. 2008) was used as a reference where all organisms were grouped into three super kingdoms namely archea, bacteria and eukarya. However, for broader understanding, all the DNMT containing organisms were grouped into seven kingdoms such as archea, bacteria, algae, fungi, plants, invertebrates and vertebrates. Each genus was classified up to phylum level and very large phyla like ascomycota/basidiomycota or angiospermata were classified up to class level. The whole classification is presented in **Tables S1, S2 and S3.** Most prevalent bacterial organisms were from phylum proteobacteria and actinobacteria. Arthropods were found to be most prevalent DNMT1 containing phyla. Most prevalent vertebrates were rodent and primate mammals; other mammals were also present in the classification. Most prevalent DNMT3A and DNMT3B containing organisms were mammals. DNMT3B is also present in plants and invertebrates. Presence of both of these enzymes only in higher order organisms suggests that they probably have originated much later than DNMT1.

### Phylogeny of DNMTs along tree of life

In order to trace the evolutionary history of DNMT enzymes along different branches of tree of life the 710 organisms were classified into seven major kingdoms as mentioned above. No archea, algae and fungi were found to possess sequences of DNMT3A and DNMT3B. Only 15 plants were detected with DNMT3B sequences but no DNMT3A sequences. Among 292 bacteria only 7 bacteria contain DNMT3A sequences and 32 genuses contain DNMT3B sequences. Similar distribution was observed in invertebrates where only one organism was detected with DNMT3A sequence and nine organisms were detected with DNMT3B sequences. On the contrary, most of the vertebrates (75%) were detected with sequences of all three DNMT enzymes. Distribution of the organisms is presented in **Table 1.** A matrix was also created to represent the number of unique organisms having any one, two or all of the three enzymes **(Figure S2)**. In 2006 Ciccarelli proposed one of the most extensive tree of life which spans across 191 genomes (Ciccarelli F. D. et. al. 2006). For detection of evolutionary lineage, 16 protein families were considered to build this huge tree. The presence of DNMT enzymes were mapped on the same tree. Only 134 organisms were common between both the dataset **(Figure 1)**.

**Figure 1:**
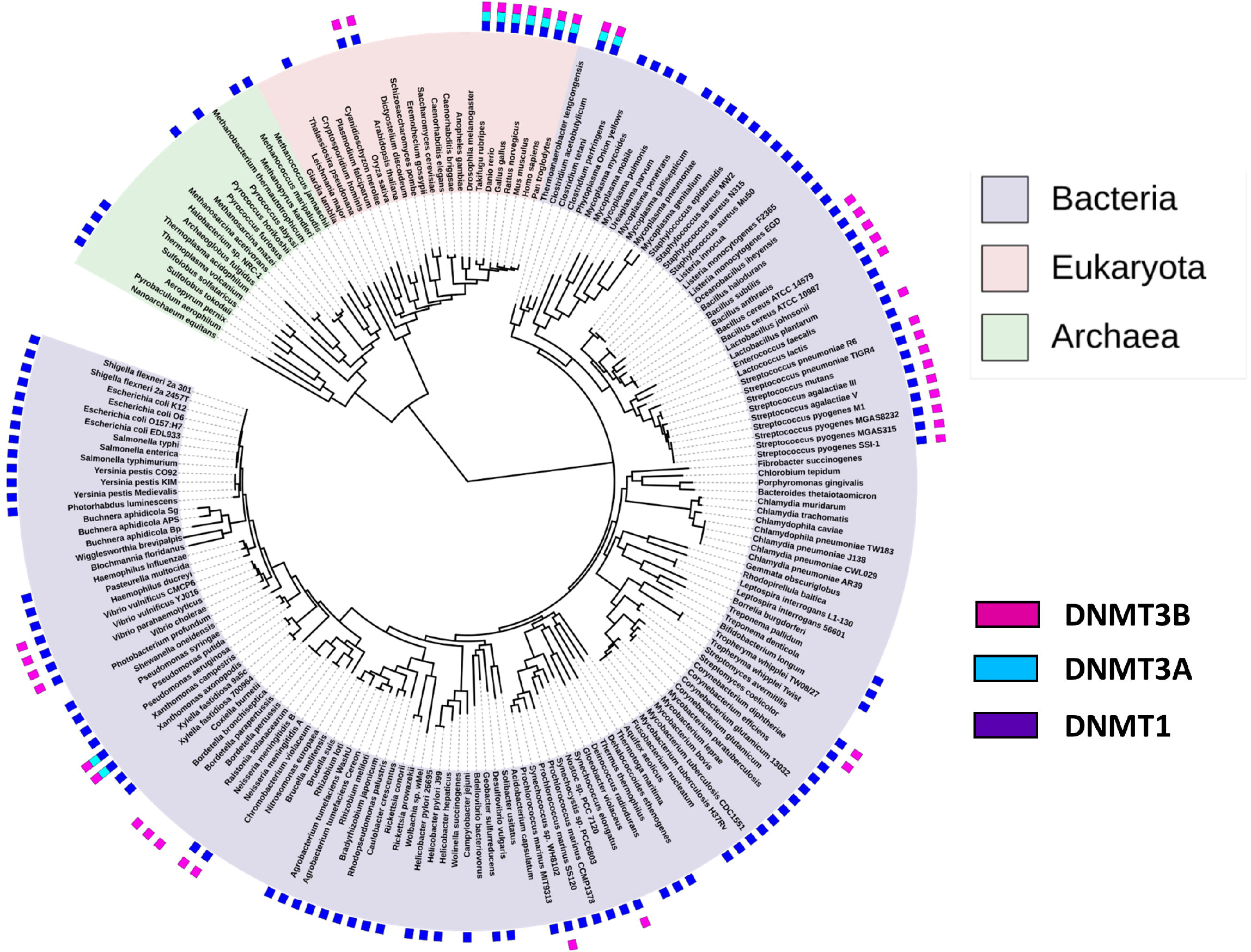
DNMTs mapped on protein based tree of life (TOL). Presence of each of the 3 DNMT enzymes (indicated by 3 different colors) is marked beside each leaf of the tree of life (Ciccarelli F. D. et. al. 2006). The tree and image was obtained from ITOL server.

**Table 1:** Number of Different DNMT containing organisms in various phylogenetic groups.

| **Kingdom level of organisms** | **Total number of organisms** | **Number of organisms containing DNMT1** | **Number of organisms containing DNMT3A** | **Number of organisms containing DNMT3B** |
| --- | --- | --- | --- | --- |
| **Bacteria** | **292** | **281** | **7** | **32** |
| **Archea** | **30** | **30** | **0** | **0** |
| **Algae** | **5** | **5** | **0** | **0** |
| **Fungi** | **46** | **46** | **0** | **0** |
| **Invertebrates** | **25** | **23** | **1** | **9** |
| **Plants** | **26** | **26** | **0** | **15** |
| **Vertebrates** | **40** | **37** | **31** | **31** |

All three DNMT sequences were subjected to multiple sequence alignment (MSA) followed by a phylogenetic tree creation. Total 448 organisms were found to contain 1773 sequences of DNMT1 sequences. Longest sequence from each organism was selected for the MSA. DNMT1 sequences were identified in all kingdoms of tree of life. In the tree, higher organism like vertebrate and invertebrates were clustered together, whereas in plants, fungi and algal species DNMT sequences are more closely related. DNMT1 in certain archea groups were more closely related to bacteria than to themselves **(Figure 2).** On the contrary DNMT3A sequences were identified only in vertebrates and bacteria. DNMT3B sequences were detected in bacteria, plants, invertebrate and vertebrates. Two important observations from this analysis were (i) unlike r-RNA based tree of life few of the bacteria and archea are closely related, (ii) eukaryotic kingdoms are well clustered among themselves and distantly related from one another.

**Figure 2:**
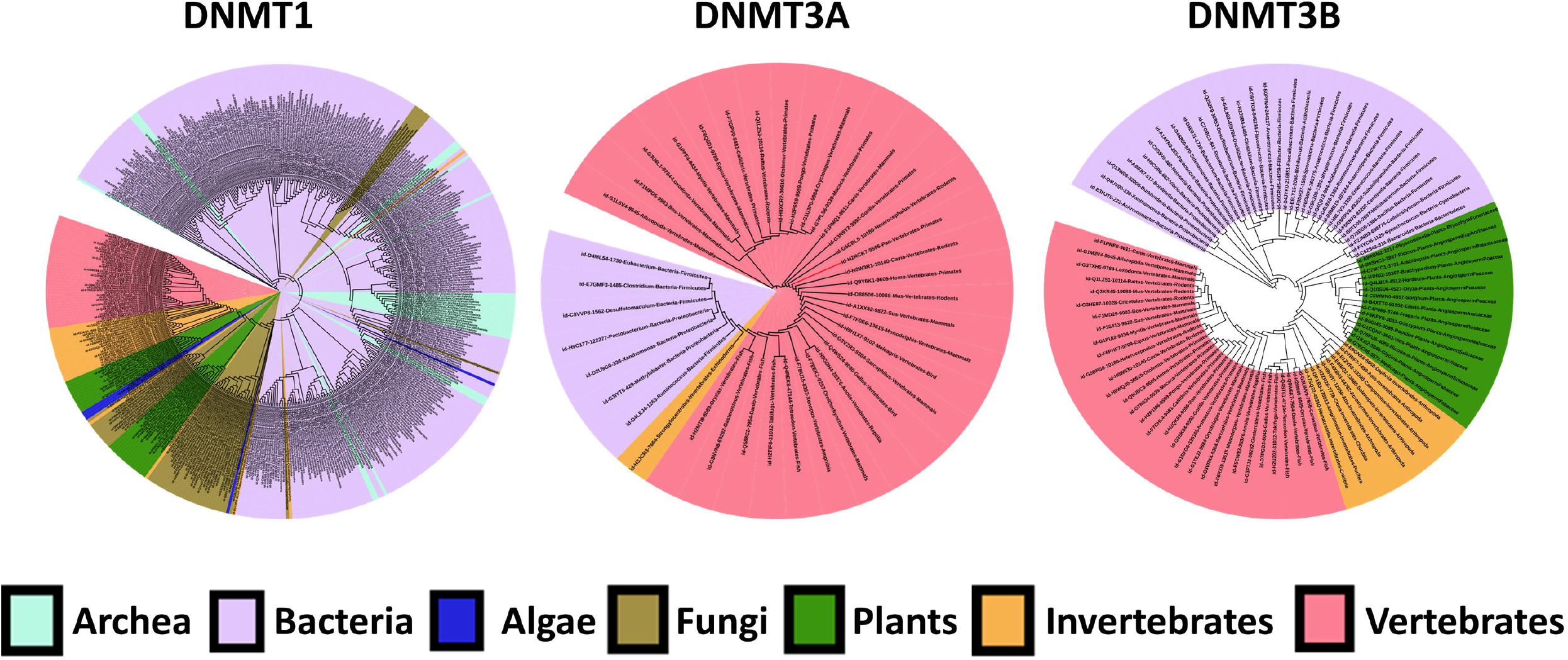
Phylogenetic trees for all DNMT1, DNMT3A and DNMT3B sequences. One full length representative sequence from each organism was used to construct the tree for each enzyme. The source organisms/leaves in the tree are color coded according to their kingdom.

### Variation in DNMT sequences among different organisms

Collection and phylogenetic analysis of the sequences were followed up by analysis of variation across seven major kingdoms of life. The variation was investigated using parameters like gene copy number, length of protein, sequence identity etc. Many organisms were identified with multiple copies of DNMT sequences. Highest copy number of DNMT1 genes (97) was found in *Helicobacter sp.,* a proteobacteria. Many other bacteria (101) were found to possess 3 or more copies of DNMT genes. *Clostridium sp.* was observed to contain as high as 28 copies of DNMT1 and 12 copies of DNMT3B. Most of the algal and fungal species were found to contain two to four copies of DNMT1 and DNMT3B. On the contrary, in higher order organism like mammals one or two copies of genes were identified. Zebra fish *Danio rerio* was observed to have highest copy number of all three enzymes having six copies of DNMT1, seven copies of DNMT3A and twelve copies DNMT3B. Few plants like *Oryza sativa* and *Vitis vinifera* were observed with DNMT1 copy number as high as 17. The distribution of copy numbers different kingdoms are presented as a box whisker plot in **Figure 3A**.

**Figure 3:**
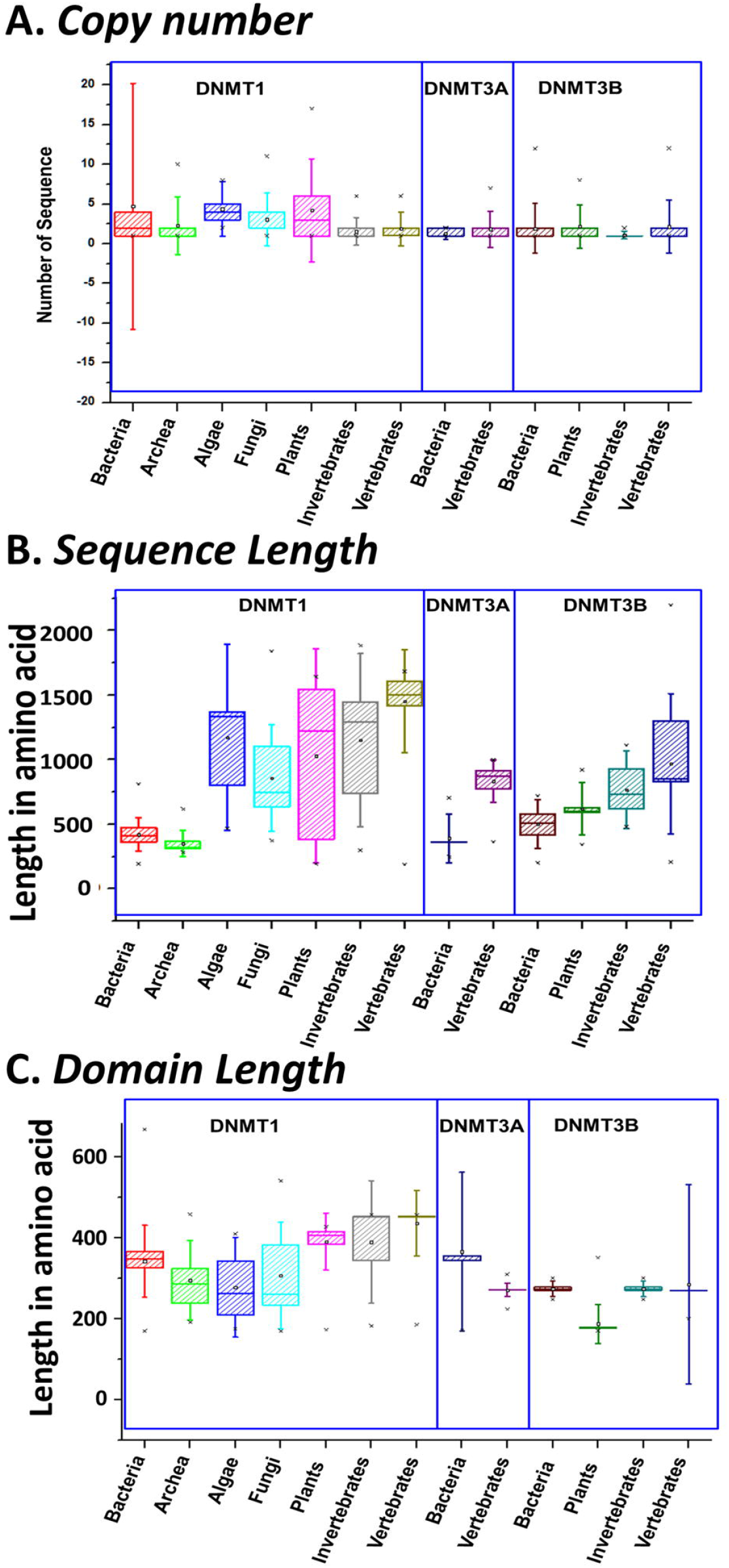
Variation in DNMT enzymes in (A) copy number (B) Sequence length and (C) Domain Length. The distributions are plotted as box whisker plot for each enzyme from each of the seven kingdoms. Boxes represent second and 3^rd^ quartile of the distribution while whiskers represent the standard deviation. Horizontal line across the box denotes the median and a small square represent the mean.

Many organisms from the same group were observed to have DNMT enzymes of different lengths. The mean length of whole enzyme increases with the evolutionary hierarchy. Average length of DNMT enzymes in bacteria is about 424 amino acids whereas algal species contain very long (>= 2000) DNMT1 sequences. Average length of vertebrate DNMT1 sequences is about 1452 amino acids while the same of DNMT3A and DNMT3B are about 834 and 970 amino acids. The distribution of sequence lengths are plotted in **Figure 3B.**

From each DNMT1 full-length sequence, Cytosine specific (C5) DNA methyltransferase domain was extracted for length comparison. Interestingly, average length of the DNMT domain in DNMT1 sequences also increases with evolutionary hierarchy while the same in DNMT3A follows the opposite trend. Average lengths of DNMT domain from DNMT3B sequences are very similar in different kingdoms. The distributions of domain lengths are presented in **Figure 3C.** Another interesting observation was that the variability in the length of whole enzyme is higher than the length of the domain.

From all the sequences the DNA methylase domains were extracted and aligned pairwise in all-to-all combinations in order to identify the overall conservation of DNA methylase domain across each group. Though functionally all of them are responsible for methyl transfer to C5 position yet the sequence identities were low in most of the groups especially in bacteria, algae and fungi. Interestingly, archea DNMT1 are relatively more conserved (51%) similar to that of plants and invertebrates. However, vertebrate DNMT1 sequences were found to be very highly conserved (80%) compared to other groups. Interestingly, both the bacterial and vertebrate DNMT3A sequences were also found to be quite highly conserved (67%) compared to DNMT3B where bacterial sequences possess significantly lower conservation pattern compared higher organisms **(Figure 4)**.

**Figure 4:**
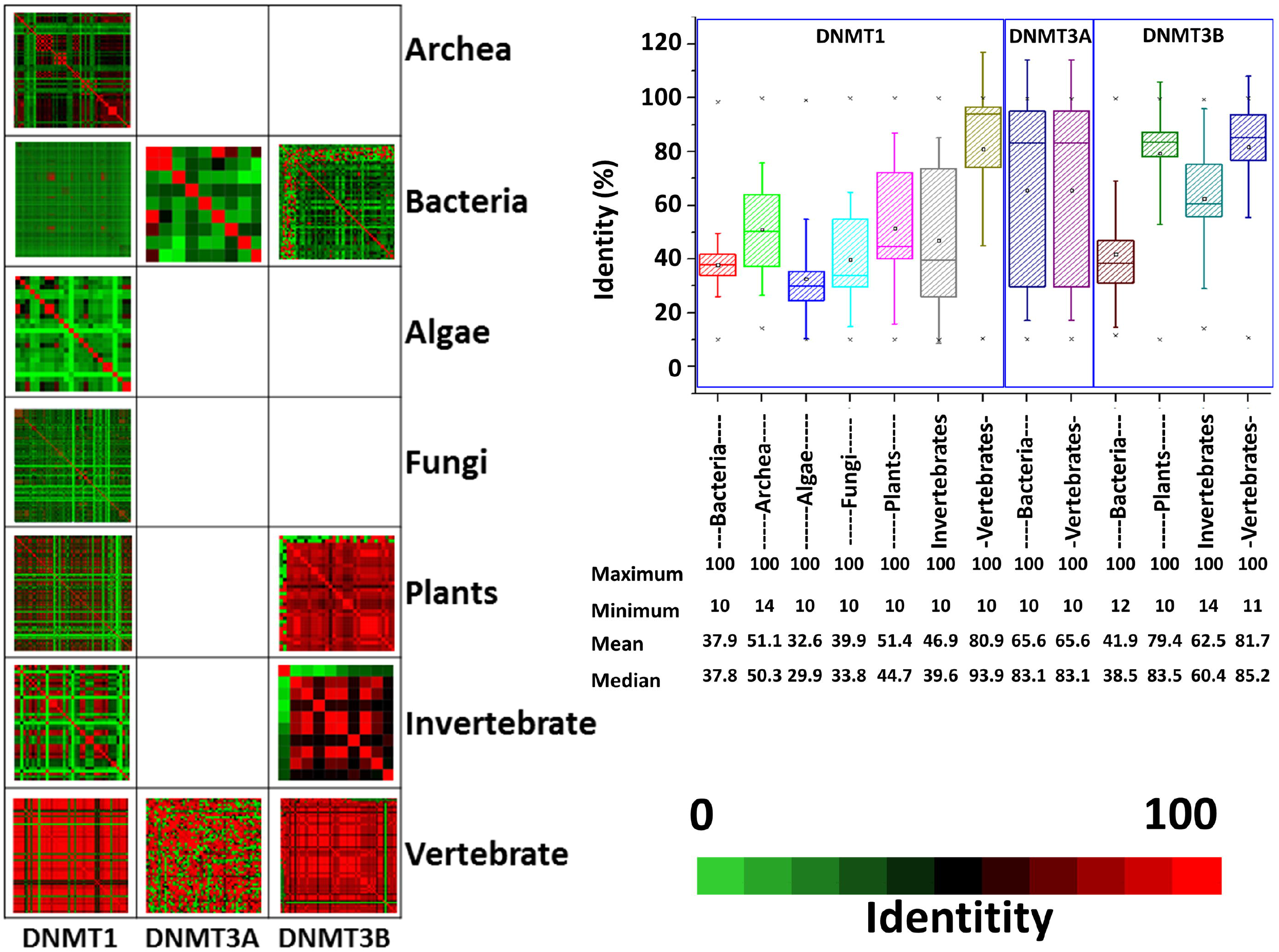
Sequence identity of DNMT enzyme across groups. In the left panel all-to-all sequence identity for each sequence in each groups for DNMT1, DNMT3A and DNMT3B are presented as color matrix and in the right panel box distributions are presented for same matrices. Percentage sequence identity is presented as a red/green color matrix. Boxes in the box plot represent second and 3^rd^ quartile of the distribution while whiskers represent the standard deviation. Horizontal line across the box denotes the median and a small square represent the mean. In the corresponding table mean and median values are presented along with maximum and minimum values of the distribution.

### Variation of DNA methyltransferase motifs and associated domains

DNA methyltransferase domain is known to have ten small motifs. All of the motifs play significant role in DNA binding, Adomet binding and catalytic activity. Motif IV (PCQ) is the catalytic motif. All these motifs were reported to be present in vertebrate DNMT1, DNMT3A and DNMT3B. As overall sequence identities were found to be low in the DNMT sequences, conservation pattern of individual motifs was examined in each of the enzyme class across each kingdom **(Figure S3)**.

All the DNMT sequences from each organism were subjected to a MSA followed by a motif scanning protocol. Though not all the motifs are present in all kingdoms and in all enzyme groups, yet six of them are found to be present in every kingdom with little or no mutation. Motifs I, which is responsible for Adomet (the methyl doner) binding, and motif IV, which is the catalytic motif, were found to be present in all organism groups. Motif VIII of DNMT3A and DNMT3B was not present in vertebrate sequences. All motifs except motif I and Motif IV are absent in DNMT3B sequences of plants. A new motif (**R**x**R**) was identified in our analysis in the DNMT1 and DNMT3A and DNMT3B sequences in all organism groups except plant DNMT3B sequences. Though the motif residues were conserved yet some changes were observed across the organisms. The glutamine (Q) in the catalytic motif IV has been replaced by asparagine (N) in many of the DNMT3A and DNMT3B sequences. Also in case of motif I (**F**x**G**x**G**) the last Glycine have been observed to have higher rate of mutation in DNMT3A and DNMT3B. Also motif I in vertebrate DNMT1 sequences (**F**S**G**C**G**) is changed into **F**D**G**I**A** in DNMT3A and DNMT3B sequences. Motif X in DNMT1 sequences (GN) is also replaced by SN in many of the DNMT3A and DNMT3B sequences. The logo plots of motifs are presented in **Figure S3** whereas an estimation of the diversity within the motifs is shown in **Figure S4**.

All DNMT sequences were scanned against Pfam database in order to map annotated protein domains in the sequences. About 275 other domains (including 61 domains with no known function) were found to be present in DNMT1 sequences but only 37 of them were present in more than 5 sequences **(Figure 5A)**. In DNMT3A sequences only one domain (PWWP) was found to be present in almost all sequences. Though 33 different domains other than DNA methylase domain were mapped on DNMT3B sequences yet only six domains were mapped onto more than 10 DNMT3B sequence.

**Figure 5:**
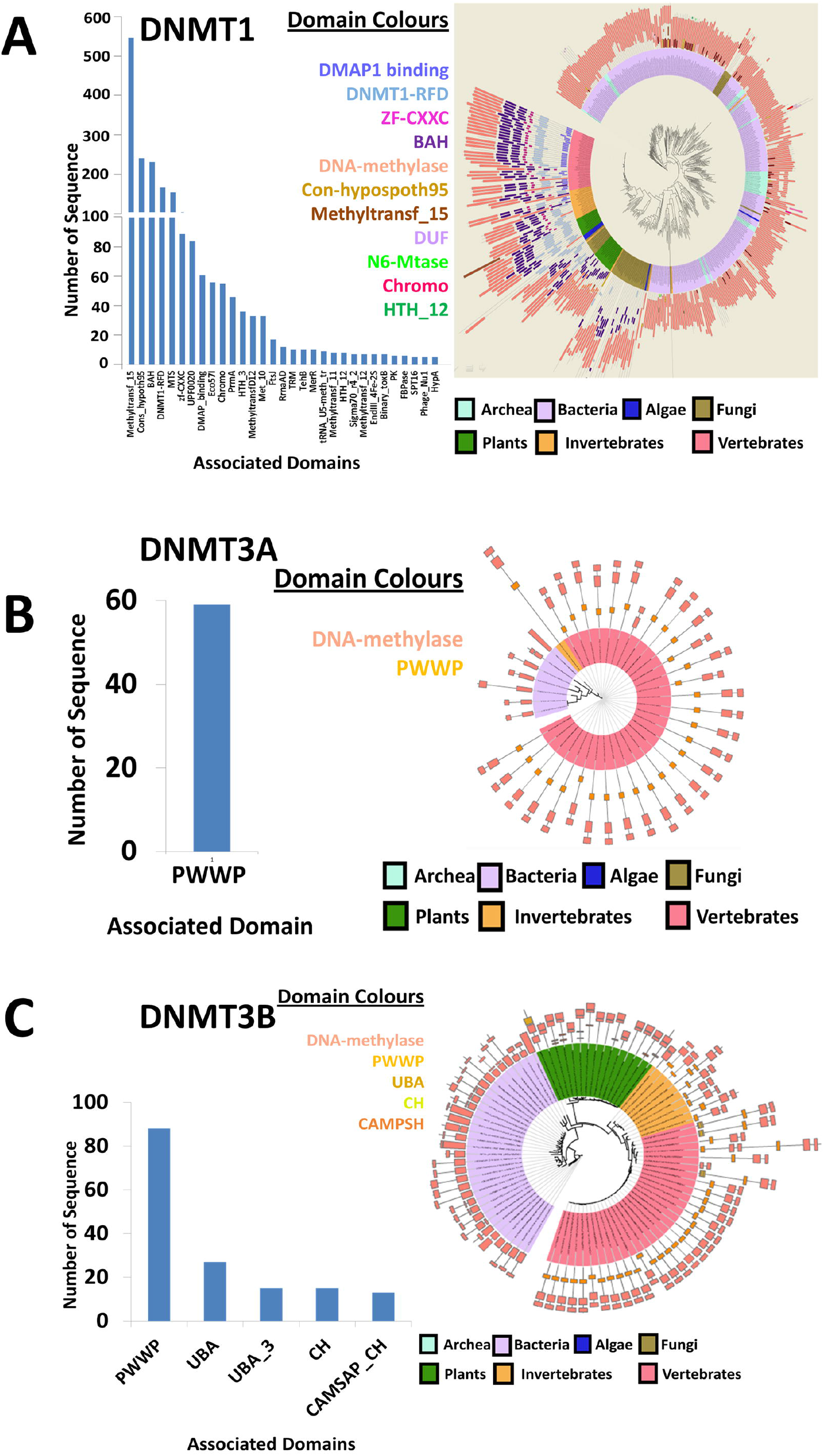
Domain architecture of DNMT1, DNMT3A and DNMT3B sequences across tree of life. In the left panel, numbers of sequences with other associated domains are presented. In the right panel the global tree of DNMT1, DNMT3A and DNMT3B are enriched with the combination of domains present in each sequence. The colors on the organisms are according to their kingdom level classification. Different domains are also marked by different colors.

All the domains were mapped onto DNMT1, DNMT3A and DNMT3B phylogenetic trees mentioned above. Almost all organism groups were observed to contain unique domain organization. Bacterial and archeal DNMT1 sequences were found to be comparatively shorter and methyltransferase domains were present at the N-terminal. Many DUF (Domain with Unknown Functions) were also mapped onto different bacterial DNMT1 sequences. Algal and fungal sequences are comparatively longer are mapped with single or double BAH domain and DNMT1-RFD domain. Fungal DNMT1 contained long stretch of sequences with no mapped domains. Interestingly, both vertebrate and invertebrate sequences were found to have multiple other domains. Many invertebrate and all vertebrate sequences were mapped with DMAP1 binding, DNMT1-RFD, zf-CXXC, two consecutive BAH domain and DNA methylase domain from N to C terminal **(Figure 5A)**. All DNMT3A sequences were mapped with only two domains PWWP at the N terminal and DNA methylase at the C-terminal **(Figure 5B)**. Most of the DNMT3B bacterial sequences were mapped with only DNA methylase domain whereas UBA domain was mapped in many plant sequences. All invertebrate and vertebrate sequences were mapped with PWWP domain. Another interesting observation was that in both DNMT3A and DNMT3B there is a large insertion inside the DNA methylase domain **(Figure 5C)**.

### Single nucleotide polymorphisms in DNMTs

A list of reported single nucleotide polymorphisms (SNPs) for all three human DNMT1, DNMT3A and DNMT3B were compiled from ensemble transcripts sequences (**Table S4**). Mutation in DNMTs could be very crucial for different diseases. In order to investigate whether the functionally important SNP positions are conserved in vertebrates, all SNPs were mapped over vertebrate specific alignment of DNMT1, DNMT3A and DNMT3B. In all three cases percentage of total SNP positions that are not conserved was found to be connected with deleterious effect, whereas most of the SNPs which are deleterious in effect were found to be more conserved **(Figure 6).** The distribution of SIFT score, which indicates the deleterious effect of a SNP and the conservation index (measured by AL2CO scores) was compared together to identify both deleterious and conserved SNPs **(Figure 7)**. Mutations map was created for all three enzymes. It was observed that most of the conserved and deleterious SNPs were result of mutation replacement of polar uncharged residues with hydrophobic residue or vice versa. On the contrary, SNPs without any deleterious effect were resulted from replacement with similar residues **(Figure 7).**

**Figure 6:**
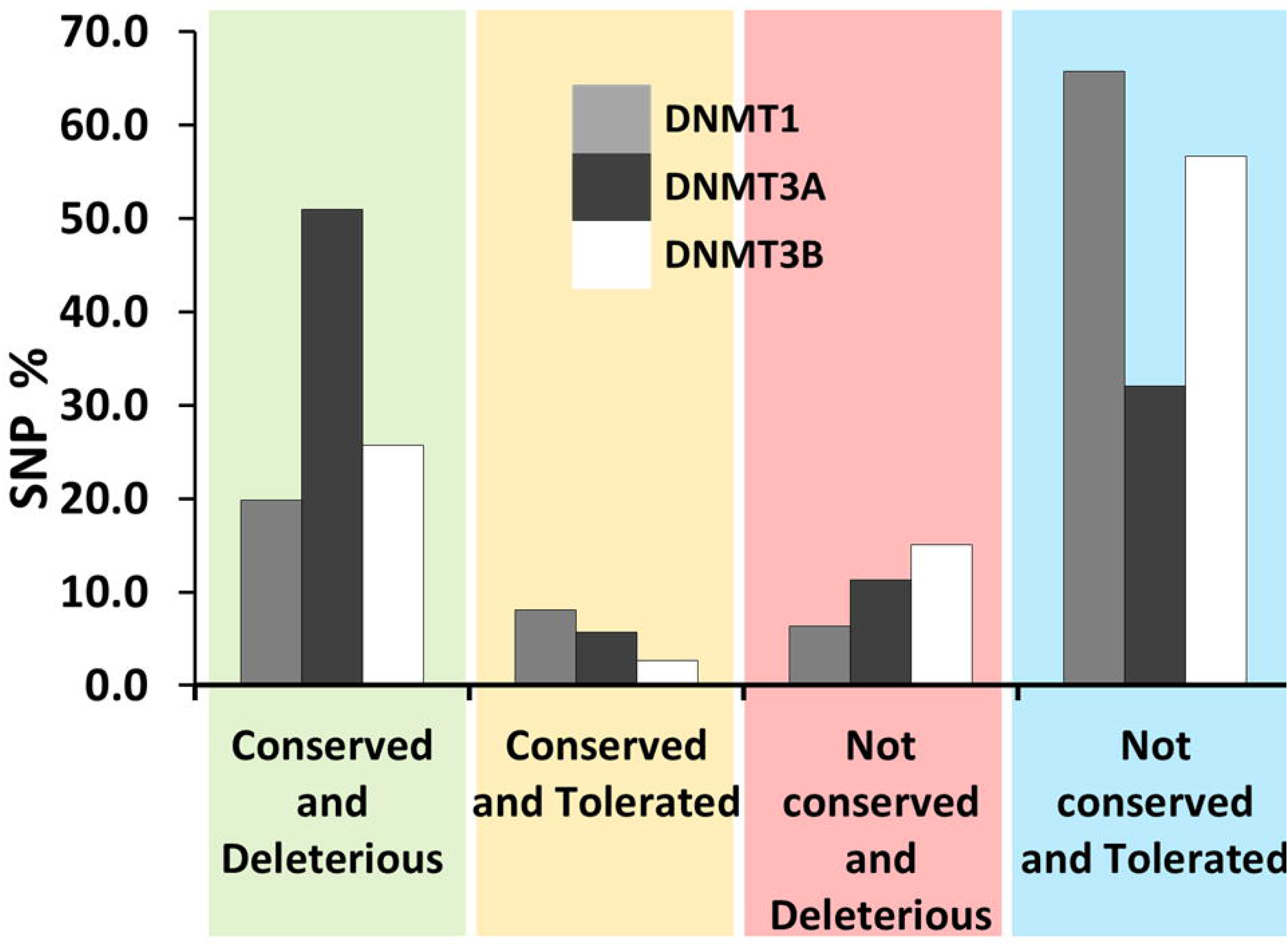
Distribution of different types of SNPs in all three DNMT enzymes. Both deleterious and tolerated SNPs are mapped and are plotted against with the sequence conservation of those particular alignment columns. Each of the 4 combinations is marked with different color as mentioned in X axis.

**Figure 7:**
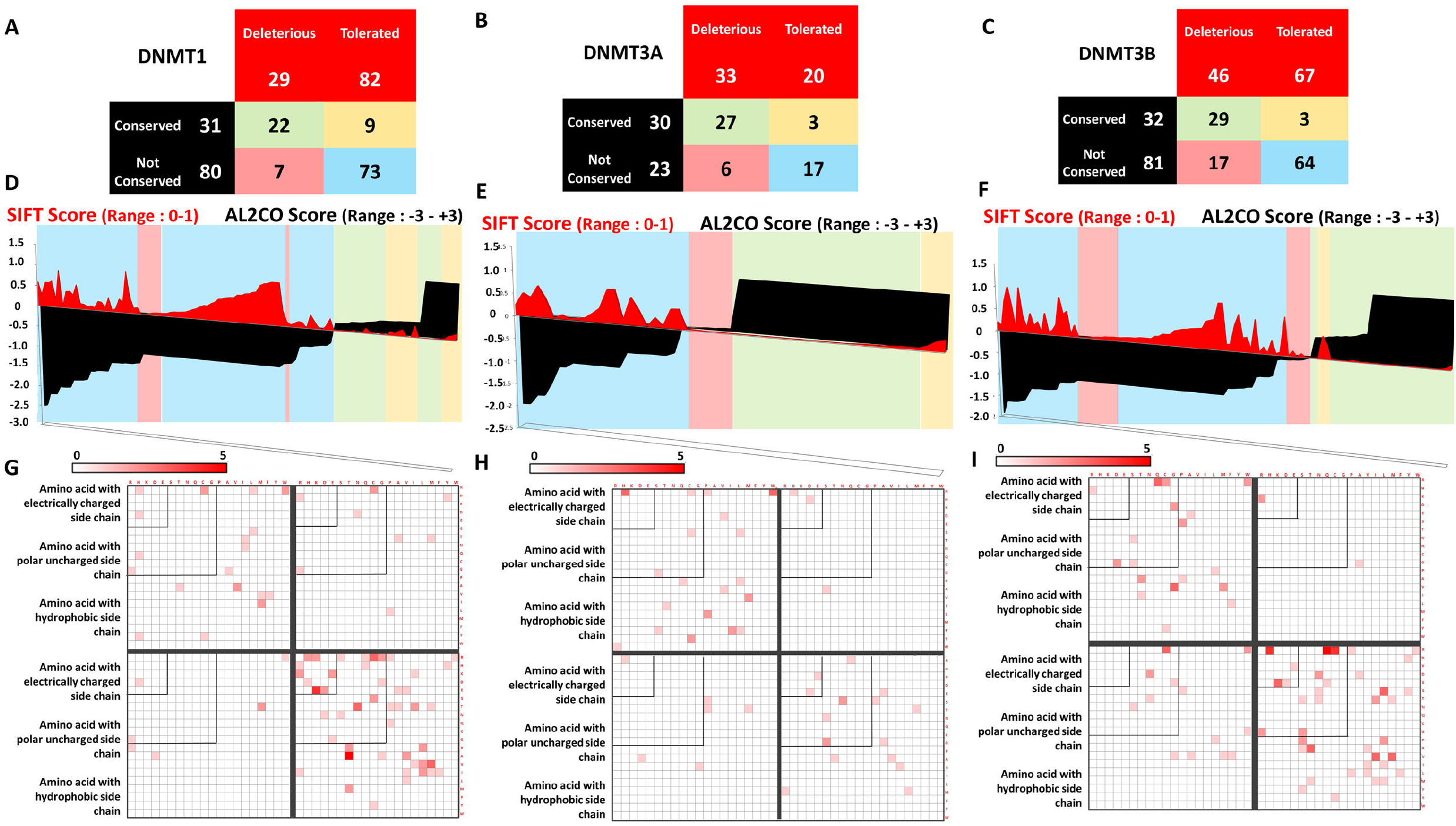
SNP conservation and mutation map for DNMT1, DNMT3A and DNMT3B. (A-C) Count of different types of SNPs based on the combination of SIFT score and Al2Co score. Each combination is indicated by a colour. (D-F) Distribution of SIFT score and Al2CO score. Background color indicates different combination range of both score as indicated in panel A. (G-I) Raw numbers of a particular mutation was marked into a 20 × 20 amino acid matrix to create a mutation map for each of the combination range of SIFT score and Al2CO score. Order of the combination is same with panel A.

## Discussion

In this study, evolutionary studies of three DNA (C5) methyl transferase enzymes namely DNMT1, DNMT3A and DNMT3B are performed. This is one of the few large scale study considering about five hundred organisms and about two thousand sequences. This was also the first attempt made towards exploring the landscape of evolutionary history of an extremely critical and important enzyme in different classes of organism. This study shows the presence of cytosine specific methyltransferase in many primitive organisms like bacteria and archea, which contradicts the common idea that cytosine specific methylation, is a regulatory mechanism present only in higher order organisms. Rather, it has been found to be a global phenomenon with conspicuous absence in few organisms like yeast, worm and fruit fly.

DNA methylation was found to play important roles in the biology of bacteria: phenomena such as timing of DNA replication, partitioning nascent chromosomes to daughter cells, repair of DNA, and timing of transposition and conjugal transfer of plasmids are sensitive to the methylation states of specific DNA regions. All of these above mentioned events use the hemimethylated state of newly replicated DNA as a signal. In the case of DNA replication, the protein SeqA binds preferentially to hemi-methylated DNA target sites (GATC sequence) clustered in the origin of replication *(oriC)* and sequesters the origin from replication initiation. In addition, SeqA also transiently blocks synthesis of the DnaA protein, which is necessary for replication initiation, by binding to hemi-methylated GATC sites in the *DnaA* promoter. In DNA repair, the methyl-directed mismatch repair protein MutH recognizes hemi-methylated DNA sites and cuts the non-methylated daughter DNA strand, ensuring that the methylated parental strand will be used as the template for repair-associated DNA synthesis. DNA methyltransferases in bacteria were best understood in the context of restriction–modification (R–M) systems, which act as bacterial immune systems against incoming DNA including phages. But several orphan methyltransferases, which were not associated with any restriction enzyme, have also been characterized and may protect against parasitism by R–M systems. Interestingly, in a report by Sandip Krishna in 2012 this orphan methyltransferases were found to be more conserved than RM methyltransferase. In our study, we have identified nearly 1400 enzymes across 450 genomes of bacteria.

Apart from bacteria, methylation mediated regulation was poorly understood in algae and fungi. A firsthand report of algal and fungal DNMTs along with their structural features and conservation pattern is provided by this study. In plants, methylation of cytosine bases was found in all sequence contexts: the symmetric CG and CHG contexts (where H is A, T, or C) and the asymmetric CHH context. Specific enzymes were reported that establish and successively maintain methylation patterns during DNA replication. It was suggested that methylation occurs predominantly at repeats and transposons (more than 90% are methylated), but approximately the 20% of genes also exhibit a certain degree of methylation. Overall, the levels of methylation in the *Arabidopsis thaliana* genome at CG, CHG, and CHH are about 24%, 6.7%, and 1.7%, respectively, but methylation within the genes is primarily restricted to CG sites and was predominantly observed in the transcribed coding region or the so called gene body (Kokus S. J, et. al 2008 and Lister R. et. al 2008). It was also reported that modestly expressed genes are more likely to be methylated within gene body, while genes expressed at high and low levels are usually less methylated. In our study it was found that DNMTs of plants acquire a certain conserved sequence with all the conserved motifs. It was also reported that Plants sequence were more closely related to the Algal and Fungal sequences than which corresponds to the protein based or RNA based tree of life (See Figure 2).

However, the sequence and structure of the enzyme has been evolved greatly from bacteria to vertebrates. Great variability has been observed in length, copy number and sequence identity of DNA (C5) methyltransferase domain when compared across kingdoms. It has been observed that the sequence conservation is greatly increased in invertebrates and vertebrates in comparison to other groups. Sequence length has been found to increase while domain lengths remain more or less conserved with the evolution indication association of other functional domains. By domain analysis it was found that vertebrates not only have a conserved methylation domain but also have other conserved domain which aid in the methyltransferase function. No such signature association of domains was observed. As a whole this study reports a history of evolution of DNA methyltransferase enzyme across different leaves of tree of life. There is a future scope to trace the origin of this enzyme if a phylogenetic analysis is performed including DNMTs and other methyltransferase enzymes like RNMT, DAM and MGMTs.

Comparison between naturally occurring SNPs and residues conserved in vertebrate alignment revealed that most of the tolerated SNPs are conserved in vertebrates. It was also observed that mutations caused by residues with similar chemical property were more tolerated than the other way round. In future the comparison of SNPs can be extended to alignment of all organisms to see similar selection pressure exists throughout all branches of tree of life.

## Methods

### Sequence collection

Sequences of DNA (C5) methyltransferase domain (PF00145) were extracted from Pfam and subjected to homology search using BLAST (Altschul S. F. et. al. 1990) against NR (Coordinator N. R. 2018) and Uniprot (Leinonen R. et. al. 2004) sequence datasets. Homologues were identified based on a threshold E-value of <= 10^-5^, query coverage >= 50% and subject coverage >= 50%.

All reviewed full length sequences of DNMT1, DNMT3A, DNMT3B and DNMTL were collected from Uniprot (Leinonen R. et. al. 2004). HMM-profile was made using MAFFT and HMMER for each of the four enzymes. Each sequence of DNMT was scanned against these profiles using *Hmmersearch* (Johnson L. S. et al. 2010)). Again the sequences were identified using a threshold of E-value <= 10^-5^, query coverage >= 50% and subject coverage>= 50%. Unique sets of homologous sequences were identified using an in house alignment scoring method.

### Obtaining the tree of life

Here, two trees of life have been used as references to detect the evolutionary relationship among DNMTs. First one was the 16S r-RNA based tree of life according to which evolution occurred in three different branches of life namely archea, bacteria and eukarya.

The second one was proposed by Ciccarelli in 2006 (Ciccarelli F. D. et. al. 2006). This tree was constructed using 236 protein families spanning over 191 genomes. With further investigation it was found that the tree has 112 unique genuses among which 71 are bacteria, 23 are eukaryotes and 18 are archea. Keeping this in mind, all the organism found to containing DNMTs were classified into seven groups; along with archea and bacteria, eukarya was classified into 5 more groups such as algae, fungi, plants, invertebrates and vertebrates.

### Phylogenetic tree construction

A common protocol was followed for construction of all phylogenetic trees in this study. Sequence set for tree construction was identified and redundancy was removed using CD-HIT at 100% (Li and Godzik 2006). Multiple sequence alignments (MSA) were created using MAFFT 5.1 (Kattoh K. et. al. 2002). Phylogenetic tree was constructed by RAXML-HPC 7.0.4 (Stamatakis A. 2006) package using maximum likelihood method (Bootstraping value was set as 100). In all seven groups DNA methylase domain was marked and extracted for tree construction. Tree images were created using ITOL (Interactive Tree of Life) server (Letunic and Borc, 2007).

### Calculation of sequence identity

Sequences from each 7 kingdom was subjected to all-to-all pairwise alignment using NEEDLE tool from EMBOSS package (Rice P. et. al. 2000) and identity matrices were created using inhouse programs. The numerical matrices were converted to colour matrices by using Matrix2png 1.0.6 tool (Pavlidis and Noble 2003).

### Scanning and identification of motifs

Sequence conservation was measured in all the kingdom level MSAs by AL2CO software package (Pei and Grishin 2001). Scores for all columns were sorted and subjected to statistical analysis. All columns with positive scores were identified as conserved column which then divided into three classes. The columns in the top quartile of the distribution were considered as highly conserved, the columns belonging to the second and third quartiles were termed as moderately conserved and the columns in last quartile were marked as conserved columns. All signature motifs were identified in conserved columns of the alignment and converted to logo plot using WEBLOGO 3.3 (Crooks G.E. et. al. 2004). The conservation score of each motif was calculated by an in-house Perl program.

### Scanning and identification of domains

All the sequences were scanned against Pfam database (Punta M. et. al. 2012) using *hmmsearch* by HMMER 2.1 (Johnson L. S. et al. 2010). Domains were identified with the threshold of Evalue 10^-5^ and subject coverage of >=50%. Position and combination of domain in each sequence were identified using in-house perl program. The domains were mapped onto the phylogenetic trees using ITOL server v2 (Letunic and Bork, 2006,).

### Collection of SNP data

Single nucleotide polymorphisms (SNPs) for all three human DNMT1, DNMT3A and DNMT3B were obtained from the Ensembl Genome Browser database (Hunt S. E. et al., 2018). This represented the pooled list of SNPs obtained from the dbSNP, ClinVar and 1000 Genomes Project that were mapped onto the Ensembl transcripts of DNMT1, DNMT3A and DNMT3B.

### Calculation of SIFT score

SIFT (Sorting Intolerant From Tolerant) uses sequence homology to predict whether an amino acid substitution will affect protein function and hence, potentially alter phenotype. For calculation of score a query protein is searched against a protein database to obtain homologous protein sequences. Sequences with appropriate sequence diversity are chosen. The chosen sequences are aligned, and for a particular position, SIFT looks at the composition of amino acids and computes the score. A SIFT score is a normalized probability of observing the new amino acid at that position, and ranges from 0 to 1 (NG and Henikoff, 2001, NG and Henikoff 2002, Sim et. al. 2012).

## List of abbreviations

DNMT: DNA methyl transferases
RNMT: RNA methyl transferase
DAM: Deoxy Adenosine Methylase
MGMT: Methylgualine (O6) DNA methyltransferase
SNP: Single Nucleotide Polymorphism
SIFT: Sorting Intolerant From Tolerant
iTOL: Interactive Tree of Life

## Declarations

### Ethics approval and consent to participate

Not applicable

### Consent for publication

### Availability of Data

Not applicable

### Authors’ contributions

MB and SC have designed computational experiments. SD has provided the experimental data. MB has performed experiments. MB, SD and SC has analyzed the results. MB, SD and SC has written the manuscript.

## Acknowledgement

SC acknowledges CSIR-IICB for infrastructural support, CSIR Network Project (BSC-0114), Systems Medicine Cluster (SyMeC) grant (GAP357) from Department of Biotechnology (DBT), High Risk High Reward Grant (HRR/2016/000093) from Department of Science and Technology (DST) for financial supports. MB acknowledges CSIR for PhD fellowship.

## Supplementary Figure Legends

**Figure S1: Identification and classification of DNMT sequences.** The identification and classification protocol along with result is described as a flow chart.

**Figure S2: Distribution of enzyme in different organism group.** Number of organisms containing unique combinations of enzymes is represented in a color matrix.

**Figure S3: Logo Plots of Motif I to X in DNMT1, DNMT3A and DNMT3B sequences across each kingdom.**

**Figure S4: Conservation Score of motifs in DNMT1, DNMT3A and DNMT3B.** Conservation score was calculated for each signature motifs from multiple sequence alignment of each kingdom for three enzymes separately. The conservation score per residue is presented as a bar diagram. Full length of the grey bar indicates 100 % conservation.

**Supplementary Table S1:** Number of DNMT1 sequences in Phylum/Class/Family/Order

**Supplementary Table S2:** Number of DNMT3A sequences in Phylum/Class/Family/Order

**Supplementary Table S3:** Number of DNMT3B sequences in Phylum/Class/Family/Order

**Supplementary Table S4:** Details of SNPs in DNMT1, DNMT3A and DNMT3B

