## Supplementary figures and images for "Origin and Evolution of DNA methyltransferases (DNMT) along the tree of life: A multi-genome survey"

### Figure S1

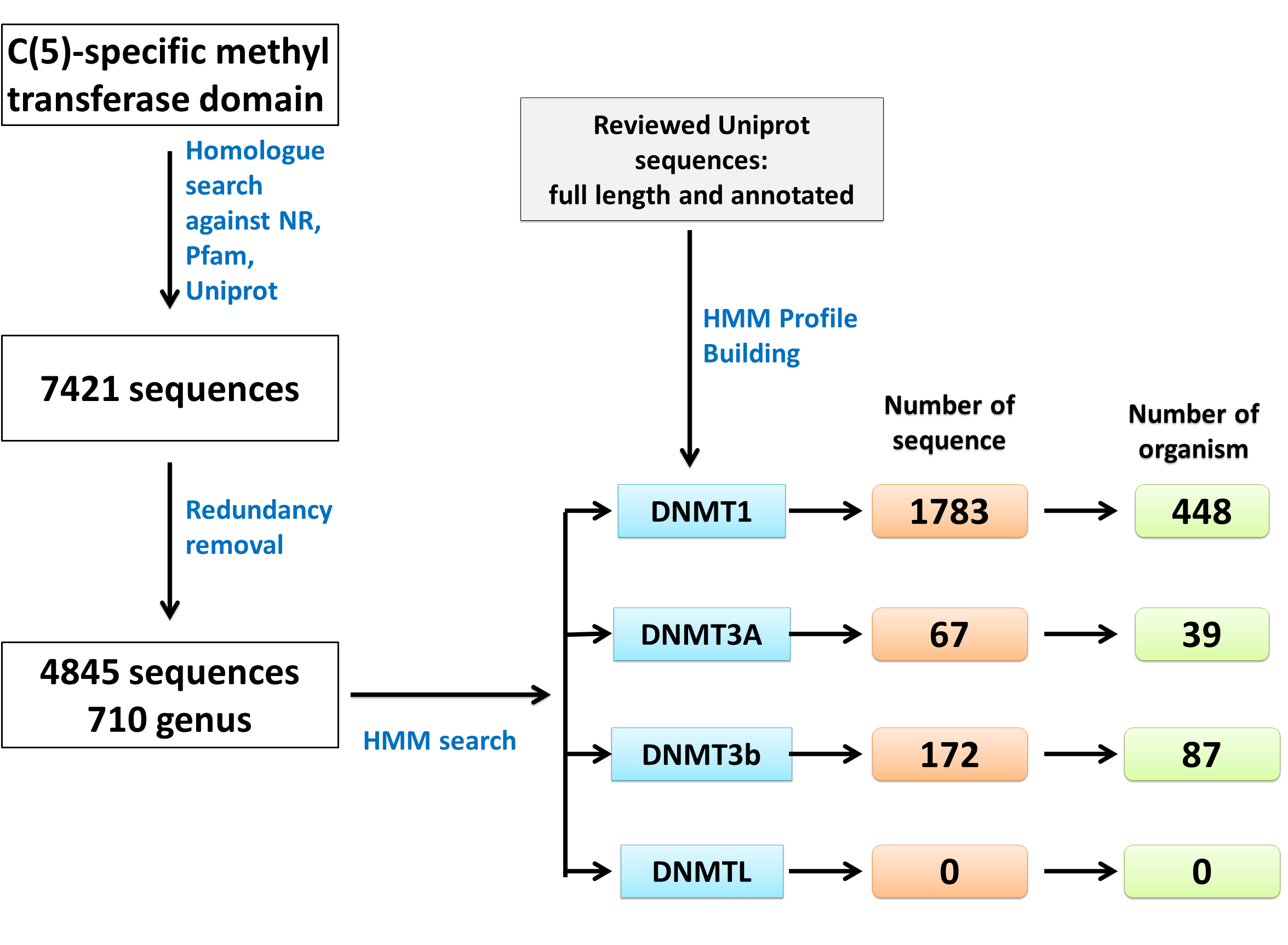

### Figure S2

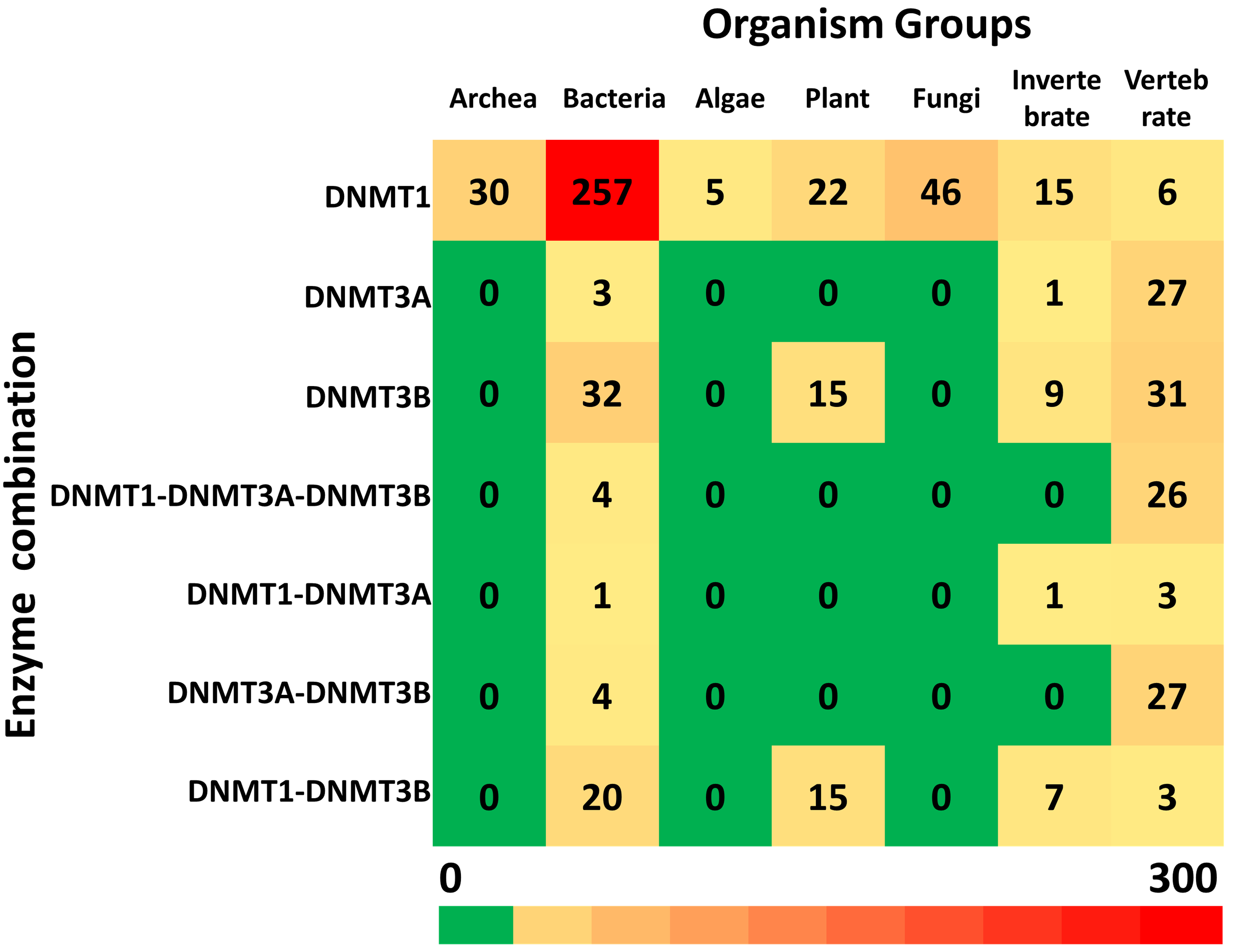

### Figure S3

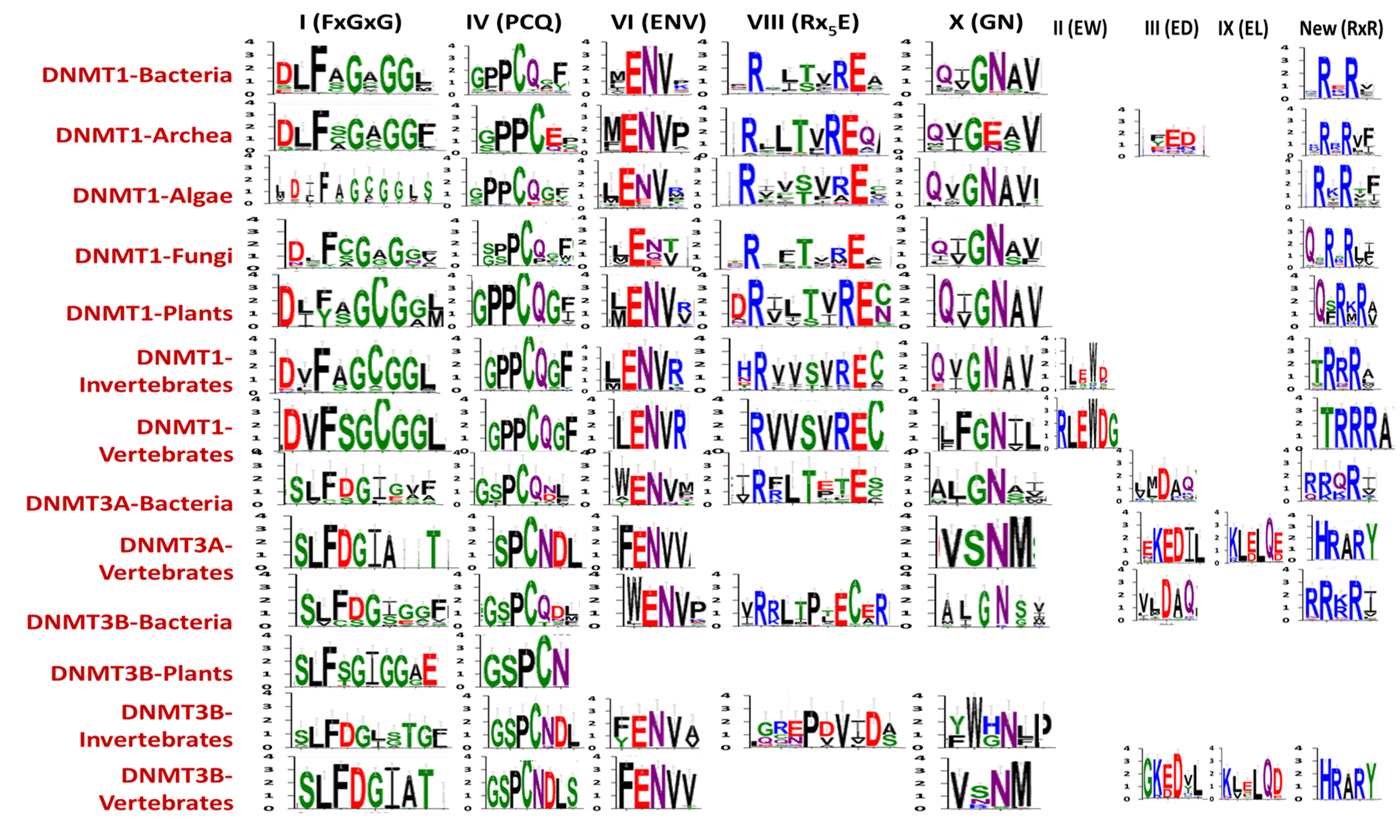

### Figure S4

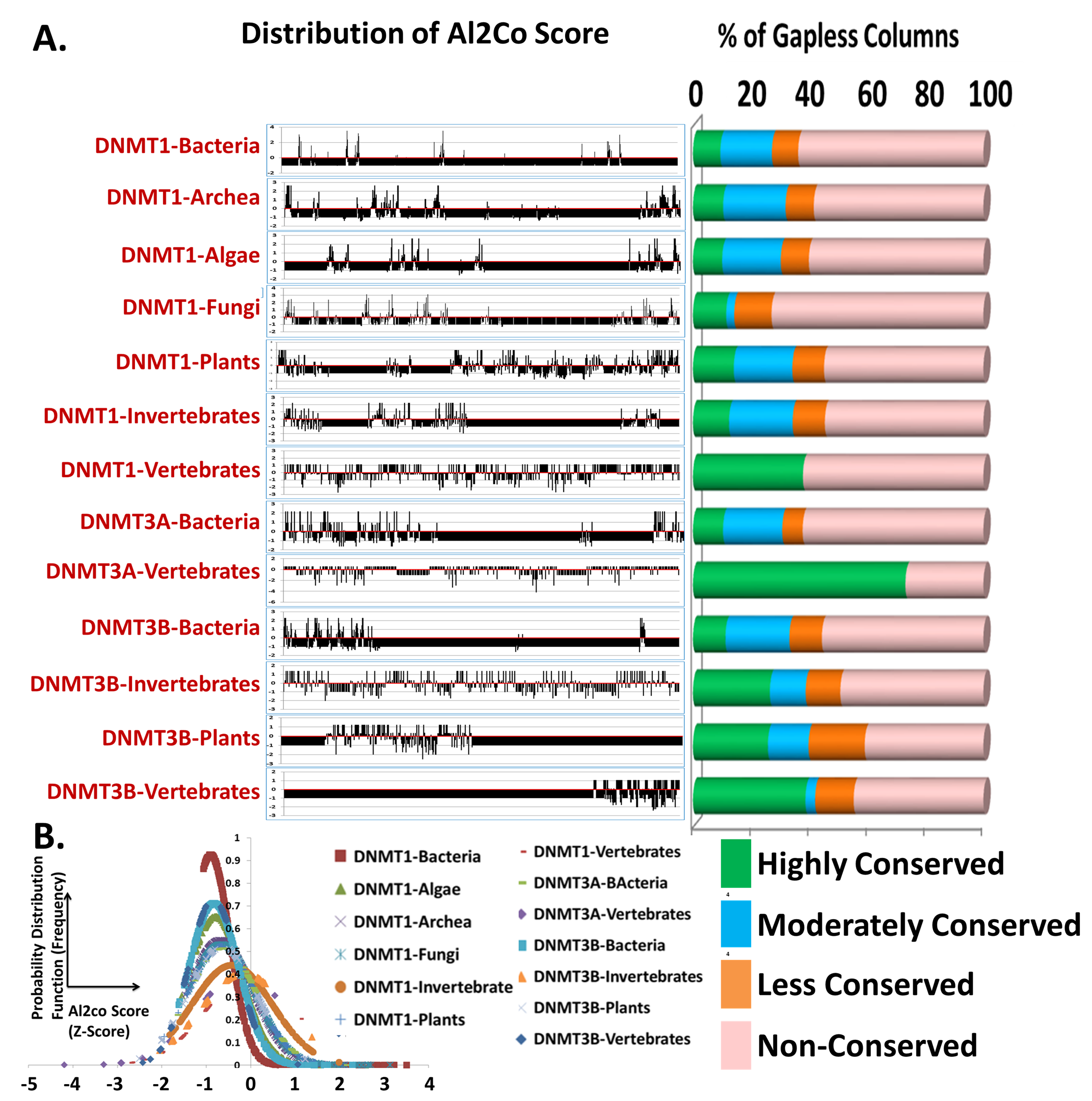
