## Supplementary material for "Origin and Evolution of DNA methyltransferases (DNMT) along the tree of life: A multi-genome survey": Table S1

Table S1: Number of DNMT1 sequences in Phylum/Class/Family/Order

| Kingdom | Phylum | Class/Family/Order |  |
| --- | --- | --- | --- |
| Algae | Chlorophyta |  | 5 |
| Archea | Crenarchaeota |  | 12 |
|  | Euryarchaeota |  | 18 |
| Bacteria | Acidobacteria |  | 2 |
|  | Actinobacteria |  | 34 |
|  | Bacteriodetes |  | 24 |
|  | Chlorobi |  | 3 |
|  | Chloroflexi |  | 3 |
|  | Cyanobacteria |  | 1 |
|  | Cyanobacteria |  | 19 |
|  | Deinococcus-Thermus |  | 2 |
|  | Fibrobacteres |  | 1 |
|  | Firmicutes |  | 51 |
|  | Fusobacteria |  | 2 |
|  | Gemmatimonadetes |  | 1 |
|  | Proteobacteria |  | 136 |
|  | Spirochaetes |  | 3 |
|  | Synergistetes |  | 1 |
|  | Tenericutes |  | 1 |
|  | Unclassified |  | 1 |
|  | Verrucomicrobia |  | 2 |
| Fungi | Ascomycota | Ajellomycetaceae | 2 |
|  |  | Arthrodermataceae | 2 |
|  |  | Ascobolaceae | 1 |
|  |  | Chaetomiaceae | 2 |
|  |  | Clavicipitaceae | 2 |
|  |  | Glomerellaceae | 1 |
|  |  | Herpotrichiellaceae | 1 |
|  |  | Hypocreaceae | 1 |
|  |  | Lasiosphaeriaceae | 1 |
|  |  | Leptosphaeriaceae | 1 |
|  |  | Magnaporthaceae | 1 |
|  |  | Mycosphaerellaceae | 1 |
|  |  | Nectriaceae | 3 |
|  |  | Onygenaceae | 2 |
|  |  | Ophiostomataceae | 1 |
|  |  | Orbiliaceae | 1 |
|  |  | Phaeosphaeriaceae | 1 |
|  |  | Plectosphaerellaceae | 1 |
|  |  | Pleosporaceae | 1 |
|  |  | Psathyrellaceae | 1 |
|  |  | Sclerotiniaceae | 2 |
|  |  | Sordariaceae | 2 |
|  |  | Trichocomaceae | 5 |
|  |  | Tuberaceae | 1 |
|  |  | Ustilaginaceae | 1 |
|  | Basidiomycota | Fomitopsidaceae | 1 |
|  |  | Marasmiaceae | 1 |
|  |  | Melampsoraceae | 1 |
|  |  | Psathyrellaceae | 1 |
|  |  | Pucciniaceae | 1 |
|  |  | Schizophyllaceae | 1 |
|  |  | Sebacinaceae | 1 |
|  | Chytridiomycota | Chytridiomycetes | 1 |
| Invertebrate | Chromalveolata |  | 2 |
|  | Annelida |  | 1 |
|  | Arthropoda |  | 14 |
|  | Chordata |  | 2 |
|  | Cnideria |  | 1 |
|  | Echinoderms |  | 2 |
|  | Hemichordata |  | 1 |
|  | Mollusca |  | 1 |
|  | Percolozoa |  | 1 |
|  | Porifera |  | 1 |
| Plant | Angiosperm | Apiaceae | 1 |
|  |  | Arecaceae | 1 |
|  |  | Asteraceae | 2 |
|  |  | Brassicaceae | 3 |
|  |  | Caryophyllaceae | 1 |
|  |  | Euphorbiaceae | 1 |
|  |  | Fabaceae | 3 |
|  |  | Malvaceae | 1 |
|  |  | Poaceae | 6 |
|  |  | Polygonaceae | 1 |
|  |  | Posidoniaceae | 1 |
|  |  | Rosaceae | 3 |
|  |  | Salicaceae | 1 |
|  |  | Solanaceae | 2 |
|  |  | Vitaceae | 1 |
|  | Bryophyta | Funariaceae | 1 |
|  |  | Marchantophyta | 1 |
|  | Pteridophyta | Lycopodiophyta | 1 |
| Vertebrates | Fish |  | 9 |
|  | Amphibia |  | 1 |
|  | Reptilia |  | 1 |
|  | Bird |  | 3 |
|  | Mammals | Primates | 8 |
|  |  | Rodents | 12 |
|  |  | Others | 5 |
