## Supplementary material for "Origin and Evolution of DNA methyltransferases (DNMT) along the tree of life: A multi-genome survey": Table S2

Table S2: Number of DNMT3A sequences in Phylum/Class/Family/Order

| Kingdom | Phylum | Class/Family/Order |  |
| --- | --- | --- | --- |
| Bacteria | Fermicutes |  | 4 |
|  | Proteobacteria |  | 3 |
| Vertebrates | Fish |  | 5 |
|  | Amphibia |  | 1 |
|  | Reptilia |  | 1 |
|  | Bird |  | 2 |
|  | Mammalia | Primates | 7 |
|  |  | Rodents | 4 |
|  |  | Others | 1 |
