## Supplementary material for "Origin and Evolution of DNA methyltransferases (DNMT) along the tree of life: A multi-genome survey": Table S3

Table S3: Number of DNMT3B sequences in Phylum/Class/Family/Order

| Kingdom | Phylum | Class/Family/Order |  |
| --- | --- | --- | --- |
| Bacteria | Actinobacteria |  | 2 |
|  | Bacteroides |  | 1 |
|  | Cyanobacteria |  | 1 |
|  | Fermicutes |  | 22 |
|  | Proteobacteria |  | 7 |
| Plantae | Angiosperm | Arecaceae | 1 |
|  |  | Brassicaceae | 1 |
|  |  | Euphorbiaceae | 1 |
|  |  | Fabacese | 2 |
|  |  | Malvaceae | 1 |
|  |  | Poaceae | 4 |
|  |  | Rosaceae | 1 |
|  |  | Salicaceae | 1 |
|  |  | Solanaceae | 1 |
|  |  | Vitaceae | 1 |
|  | Bryophyta | Funariaceae | 1 |
| Invertebrate | Porifera |  | 1 |
|  | Cnidaria |  | 1 |
|  | Arthropoda |  | 6 |
|  | Chordata |  | 1 |
| Vertebrates | Fish |  | 7 |
|  | Reptilia |  | 1 |
|  | Mammalia | Primates | 8 |
|  |  | Rodents | 5 |
|  |  | Others | 10 |
